# Detecting Somatic Mutations in Rare Clones using Single Cell Multi-Omics

**DOI:** 10.1101/2025.10.30.685685

**Authors:** Rhys Gillman, Sonam Dudka, Jerome Samir, Raymond Louie, Chris Goodnow, Fabio Luciano, Mandeep Singh, Matt A. Field

**Affiliations:** Centre for Tropical Bioinformatics and Molecular Biology, College Science and Engineering, James Cook University, Cairns, Queensland, Australia; School of Computer Science and Engineering, UNSW Sydney; Sydney, NSW, Australia; Immunogenomics Lab, Garvan Institute of Medical Research, Darlinghurst, NSW, Australia

## Abstract

**Background:** Somatic mutations are increasingly recognised as drivers of diseases beyond cancer, including autoimmune disorders. However, identifying rare, cell type-specific causal mutations remains challenging due to their low frequency within heterogeneous cell populations. Traditional bulk sequencing approaches lack the resolution to detect rare variants, underscoring the need for novel methods specifically designed for single-cell data of heterogeneous cell populations.

**Methods:** We present an integrated single-cell multi-omics computational framework, SCARCE (**S**ingle-**C**ell **A**nalysis of **R**are **C**lonal **E**vents), tailored for single-cell DNA sequencing (scDNA-seq) to statistically prioritise rare somatic mutations within defined cell subpopulations. By comparing variant frequencies across subpopulations, identified through either variant-based clustering or cell type annotation from surface marker expression, we identify variants enriched in specific cell populations. Our method applies multiple user-adjustable filters and statistical enrichment tests to distinguish true somatic variants from technical artifacts.

**Results:** SCARCE successfully identifies rare somatic mutations across three distinct datasets using technologies including MissionBio Tapestri and clonally-amplified whole-genome sequencing of single cells. We demonstrate that SCARCE correctly isolates and identifies true variants in a cell population comprising just 10 of 16,316 cells (0.06% of the total population). Furthermore, in an extensively characterised sample with known causal variants, SCARCE correctly identifies all known pathogenic variants among its top-ranked candidates.

**Conclusions:** SCARCE offers several advantages over existing tools in the field. By integrating genetic and phenotypic information at single-cell resolution, our approach opens new avenues for understanding the clonal origins of diseases driven by somatic mutations in small cell populations.

## Introduction

Somatic mutations are increasingly recognised as driving a wide range of diseases beyond cancer (Solis-Moruno et al. 2023). Neurological diseases can be driven by somatic mutations, with mutations in genes like *MTOR* and *AKT3* causing focal cortical dysplasia and epilepsy (Lim et al. 2015). Age-related diseases also exhibit somatic mutation signatures, including clonal haematopoiesis of indeterminate potential (CHIP), where acquired mutations in hematopoietic stem cells drive cardiovascular disease and increased mortality risk independent of traditional cancer development (Libby et al. 2019). Additionally, somatic mutations accumulate in various tissues during aging, contributing to organ dysfunction and age-related pathologies such as atherosclerosis, where mutations in vascular cells promote inflammatory responses and plaque formation (Andreassi et al. 2000). Finally, in autoimmune and inflammatory disorders, somatic mutations can create pathogenic self-reactive cell clones that escape normal negative selection, leading to conditions such as autoimmune lymphoproliferative syndrome, where mutations in apoptosis-regulating genes cause aberrant lymphocyte survival and autoimmunity (Holzelova et al. 2004). While somatic mutations in cancer typically drive uncontrolled proliferation, non-cancer somatic driver mutations often result in more subtle functional alterations affecting cell behaviour, developmental processes or immune regulation.

Bulk sequencing methods such as whole-genome (WGS) or exome sequencing (WES) have been employed successfully to identify germline driver mutations in a wide range of complex diseases (Field 2020; Hamzeh et al. 2021; Field 2022; Dukda et al. 2025) and cancers (Morin et al. 2011; Wilmott et al. 2015). Generally, however, bulk methods have proven unsuitable for detecting rare somatic mutations in mixed cell populations due to their inherent limitations in two critical areas: i) sensitivity and resolution of the technology, ii) the inability to differentiate cell types. Particularly challenging are cases where a somatic mutation occurs in a small subset of cells, such as in clonal haematopoiesis or early-stage cancer. These rare mutations may be present at very low allele frequencies within the heterogeneous population of cells. Since bulk sequencing captures a mixture of DNA from heterogeneous cell populations, the rare mutant alleles can be masked by the more abundant wild-type alleles, making it difficult to detect such mutations. Further, the sensitivity of bulk sequencing is often not high enough to distinguish these mutations from sequencing errors or background noise, leading to high false negative rates (Field et al. 2019). In contrast, techniques like deep sequencing or single-cell sequencing are better suited for detecting rare somatic mutations, as they offer higher sensitivity by sequencing more reads and/or analyzing individual cells, which increases the likelihood of capturing mutations present in a small fraction of the population (Dou et al. 2018).

Early methods to identify rare, cell type-specific somatic mutations employed molecule tagging using a unique molecular identifier (UMI) to input DNA either prior to or simultaneously with amplification of target sequences. Processing such data required UMI aware variant detection software such as UMI-VarCal (Sater et al. 2020) or DeepSNVMiner (Andrews et al. 2016). However, these tools often fail to scale to large numbers of samples. More recently, advances in molecular barcoding technology have led to the development of an increasing number of single-cell platforms capable of examining thousands of cells independently. These increasingly multi-omic platforms are promising for a wide variety of applications; however, most have primarily focused on RNA-based applications which are known to be problematic for variant detection for a variety of reasons including incomplete transcript representation, isoform complexity and expression of mutated alleles (Lahnemann et al. 2020). Early multi-omic platforms such as G&T-Seq enabled single cell DNA-based variant calling, where cells could be pre-sorted into disease and control groups for somatic variant detection on a small numbers of cells (Singh et al. 2020a; Young et al. 2025). More recently, high throughput single cell DNA based platforms such as MissionBio Tapestri can profile thousands of cells to identify increasingly rare somatic mutations; however challenges in differentiating cell types persist for some platforms (Macaulay et al. 2015).

To differentiate cell types, single-cell protein sequencing of cell surface markers represents a powerful approach for high-resolution cell phenotyping, which has revolutionised our understanding of cellular heterogeneity within complex tissues. This technology, primarily implemented through methods such as Cellular Indexing of Transcriptomes and Epitopes by sequencing (CITE-seq), AbSeq, and related antibody-based approaches, known collectively as antibody-derived tags (ADT), enables simultaneous measurement of dozens to hundreds of surface proteins at single-cell resolution (Stoeckius et al. 2017). While currently largely employed in differentiating immune cell types using combinations of well-known markers, cell surface markers are increasingly being employed in other applications including solid tissue biopsies (Lischetti et al. 2023), greatly broadening its application. When simultaneously combined with single cell DNA-based technologies, this approach enables the comparison of genotypes between specific cell populations, providing a powerful approach for identifying cell type-specific somatic mutations (Singh et al. 2025).

Despite these advances, there are currently limited options for simultaneous end-to-end processing of these single-cell DNA sequencing and ADT datasets. For example, several options are available for Tapestri data including Optima (Pei et al. 2023) and Mosaic (Zhang et al. 2023) packages, however, these tools are primarily limited to filtering and visualization functions. To identify such mutations, we developed an R package, SCARCE (**S**ingle-**C**ell **A**nalysis of **R**are **C**lonal **E**vents), for statistical prioritization of enriched variants in rare cell populations. When ADT data are generated, cell types are annotated in a semi-automated manner, providing the user with various visualizations to decide on final cell type annotations. Variants are annotated using VEP (McLaren et al. 2010) and filters can be applied at three levels: per variant call, within cell types, and across all cells. The workflow prioritises variants enriched within each cell type, making the identification of cell type-specific somatic mutations simple and intuitive.

## Results

We first compared SCARCE’s somatic variant enrichment (**Figure 1**) with Mission Bio’s in-house tools Mosaic and the R package Optima.

**Figure 1:**
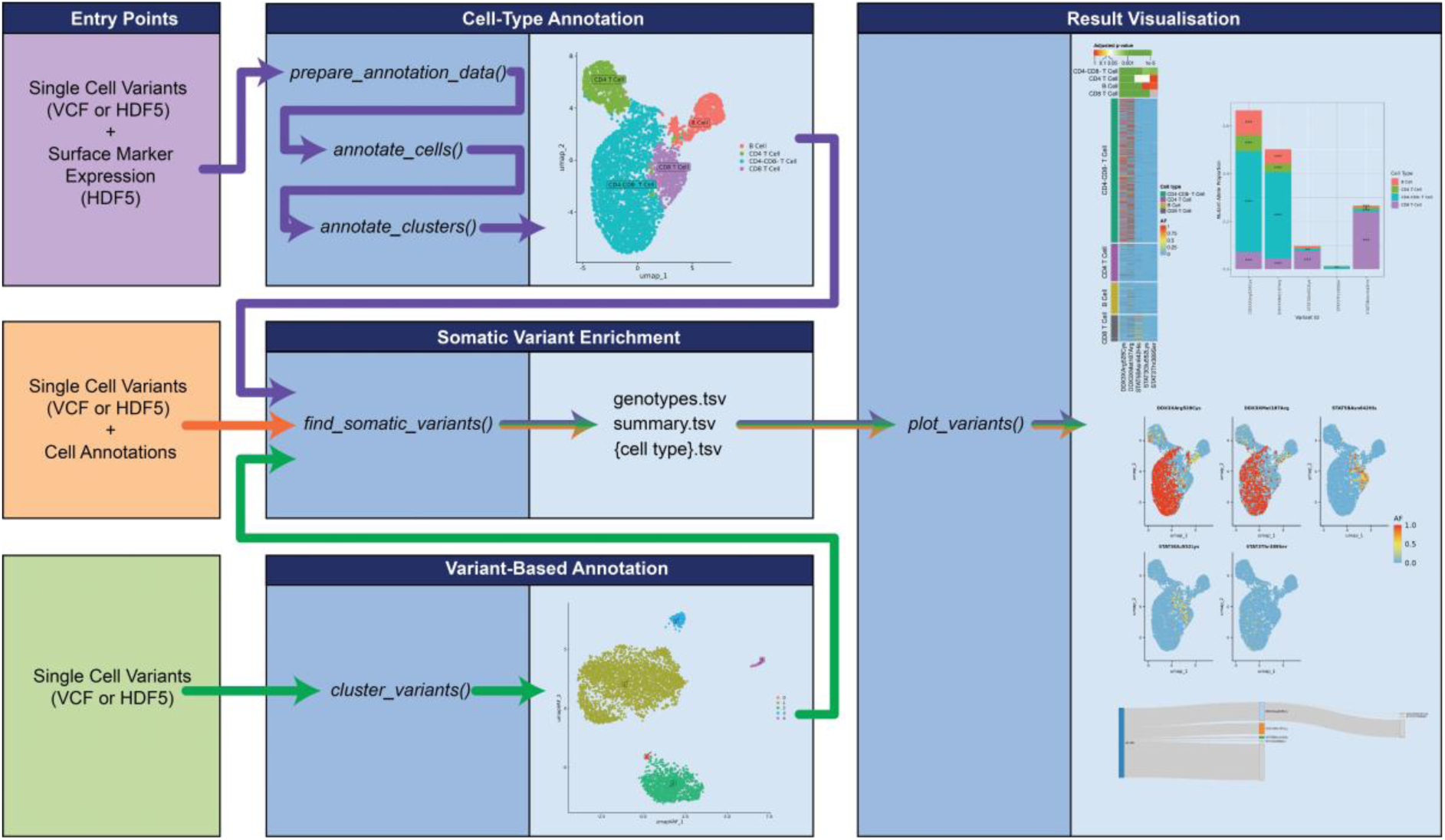
SCARCE Workflow diagram. SCARCE offers three different workflows depending on the input data available. The single-cell variants with surface marker expression pathway (purple path) begins with semi-automated cell type annotation based on ADT surface marker expression to define cell clusters. Users may alternatively provide their own cell type annotations to skip this step (orange path). Alternatively, if no surface marker expression data are available, cells can instead be clustered based on their variant genotypes to identify clusters carrying unique somatic variants (green path). All pathways converge at somatic variant enrichment, which statistically prioritises variants enriched in cell clusters, followed by visualisation of these results.

Each tool was used to analyse Tapestri scDNA-seq data from an example dataset retrieved from the Mission Bio public data repository. We analysed data from 16,316 cells with 114,788 raw variant calls derived from a mixed sample containing two peripheral blood mononuclear cells (PBMCs) samples spiked with 0.06% cells of the KG-1 acute myeloid leukaemia macrophage cell line. The targeted sequencing panel employed was the Tapestri Single-Cell DNA Myeloid panel. We sought to determine whether the tools could successfully identify variants unique to the KG-1 cells, which were expected to be represented by approximately 10 or fewer cells total. This would serve as a proxy and an extreme test case for the lower limits for somatic variant detection.

Both Optima and Mosaic offer cell-clustering based on variant genotypes, so for comparison we used the *SCARCE::cluster_variants()* function to perform the same task using the variant-only workflow. Cell populations that have acquired somatic mutations are expected to form independent clusters. As expected, SCARCE identified three clusters, including two of approximately equal size, likely corresponding to the two PBMC samples, and a small cluster consisting of only 10 cells (**Figure 2A**). Neither Optima or Mosaic, which cluster cells based on a much smaller number of variants, appeared to successfully isolate a cluster corresponding with the KG-1 cells (**Figure 2B-C**).

**Figure 2:**
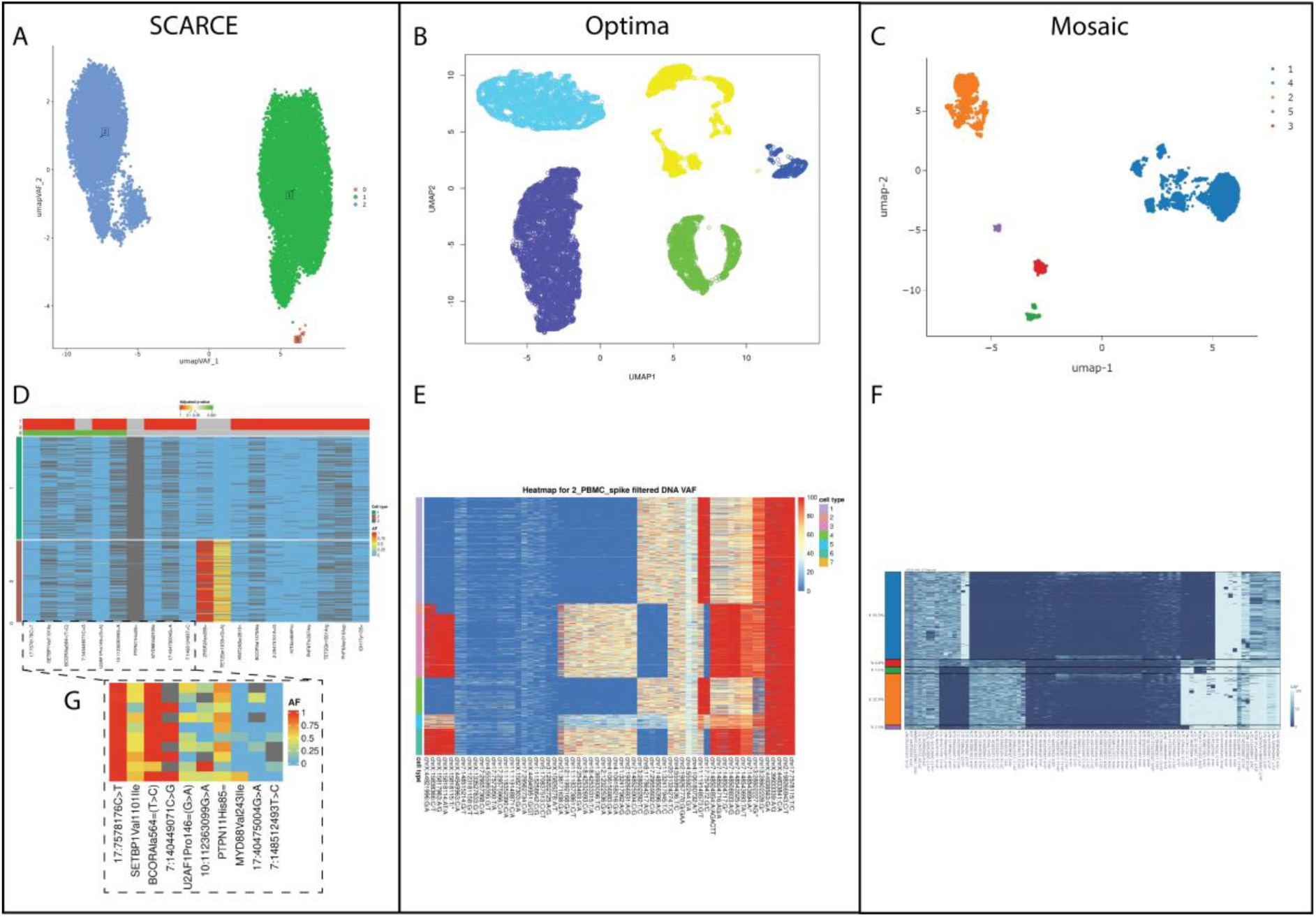
**Somatic variant detection comparison of SCARCE with Optima and Mosaic**. Tapestri sequencing data from 16,316 cells were analysed from a sample composed of a 50:50 mix of PBMCs with a 0.05% spike-in of the KG-1 cell line. Variant-based cell clustering was performed by SCARCE (A), Optima (B) and Mosaic (C). Heatmaps were also generated to visualise variant enrichment in the cell type clusters using each tool, respectively (D-F). For SCARCE, variants are ordered by p-value (Wald p, odds ratio (OR)) and filtered to p<0.05, OR > 5. An expanded view of the top variants identified by SCARCE showing just the cells for cluster 0 is also shown (G).

All three tools support generating heatmaps of filtered variants which can help visualise variant enrichment in a cell population (**Figure 2E-F**). Notably, however, these tools do not offer statistical methods for variant enrichment or prioritisation. While several variants appeared to be significantly depleted in these small cell clusters, no obviously enriched variants were observed in the output of these tools.

The ability to prioritise variants based on enrichment allows SCARCE to be much less stringent with its initial filtering of variants, allowing the detection of real somatic variants that may be filtered out by other tools. The heatmap produced by *SCARCE::plot_variants()* (**Figure 2D**) displays a p-value for variant enrichment in each cell population based on a Wald test for the mutant allele odds ratio (OR). Additionally, variants are ordered by these p-values for prioritisation and may be filtered to display only significantly enriched variants or using thresholds for OR or Wald z-score. Using SCARCE, we generated a heatmap of the top 20 variants predicted to be somatic in cluster 0, which revealed 6 variants with significant (p < 0.05) enrichment. Notably, none of these 6 variants passed the stringent filtering applied by Optima and Mosaic, and were therefore not included in their heatmaps (**Figure 2E-F**).

To determine whether the variants identified by SCARCE were true positives, we retrieved a list of known variants in the KG-1 cell line from Depmap (https://depmap.org/portal/). Due to the targeted sequencing used in the Tapestri platform, only a single variant overlapped with a known variant in the cell line: a non-coding splice-donor variant in *TP53* (build hg37, chr17:7578176C>T). This variant was identified by SCARCE and ranked as the most highly enriched variant. Given that this variant was identified in 10/10 (100%) cells in cluster 0 and in only 30/15,734 (0.2%) of the other clusters, we confirmed cluster 0 as the KG-1 cell line.

Additionally, SCARCE offers several additional visualisations to help users identify somatic variants. **Figure 3A** shows a stacked bar plot of mutant allele proportions per cell-type with significant enrichment highlighted. The distribution of called genotypes or allele frequencies can also be visualised on a UMAP (**Figure 3B**). Finally, SCARCE can produce two plots that assist in the identification of variants shared between multiple cells, an UpSet plot (**Figure 3C**) and a Sankey plot (**Figure 3D**). These can be particularly useful for identifying clonal driver mutations and clonal lineages.

**Figure 3:**
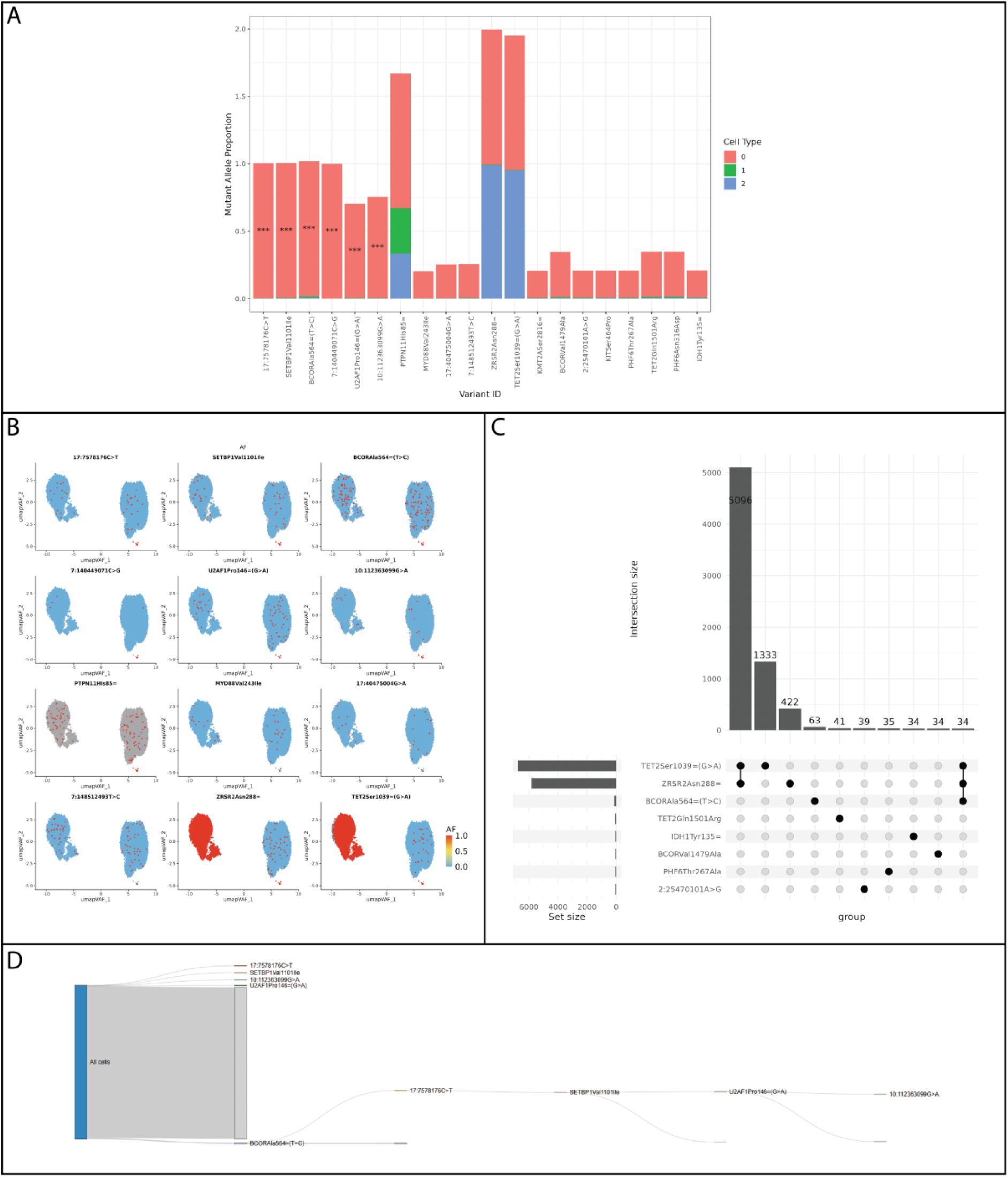
**Additional visualisations of top variants prioritised by SCARCE**. A) A stacked bar plot indicating mutant allele proportions in each cell type, with enrichment significance (Wald p, OR) indicated for each cell type. B) UMAP plots coloured by mutant allele frequency per cell of prioritised variants. C) UpSet plot illustrating set sizes and set overlaps of called variants to indicate variant co-occurrence. D) Sankey plot, indicating proportions of cells carrying significantly enriched variants and the presence of co-occurring variant sets.

Next, we ran all three tools to identify driver somatic variants in lymphocytes from a patient with refractory coeliac disease (RCD) using Tapestri sequencing data for 4,858 cells. Importantly, this sample was previously characterised extensively using multiple sequencing technologies to identify five likely driver mutations: *DDX3X^p.M187R^*, *DDX3X^p.R528C^*, *SOCS3^p.V84I^*, *STAT5B^p.N642H^*, *STAT3^p.E552K^* (Singh et al. 2025). All three tools returned four of the five driver mutations in their filtered results except for *SOCS3^p.V84I^*, which was filtered by all three tools due to low depth reads and poor quality alignment scores. **Figures 4B-C** show the heatmaps produced by Optima and Mosaic, respectively. In the cell clusters produced, only two of these driver variants show obvious sub-population enrichment, namely *DDX3X^p.R528C^* and *DDX3X^p.M187R^*. In contrast, the somatic variant prioritisation performed by SCARCE identifies all four of these variants that passed filtering with high significance, and two additional potential somatic variants are identified: *STAT3^Ser614Arg^*and *STAT3^Thr389Ser^* (**Figure 4A**). The UMAP visualises the enrichment of these variants, and the Sankey plot (**Figure 4E**) again shows limited co-occurrence, in agreement with the previous findings (Singh et al. 2025).

**Figure 4:**
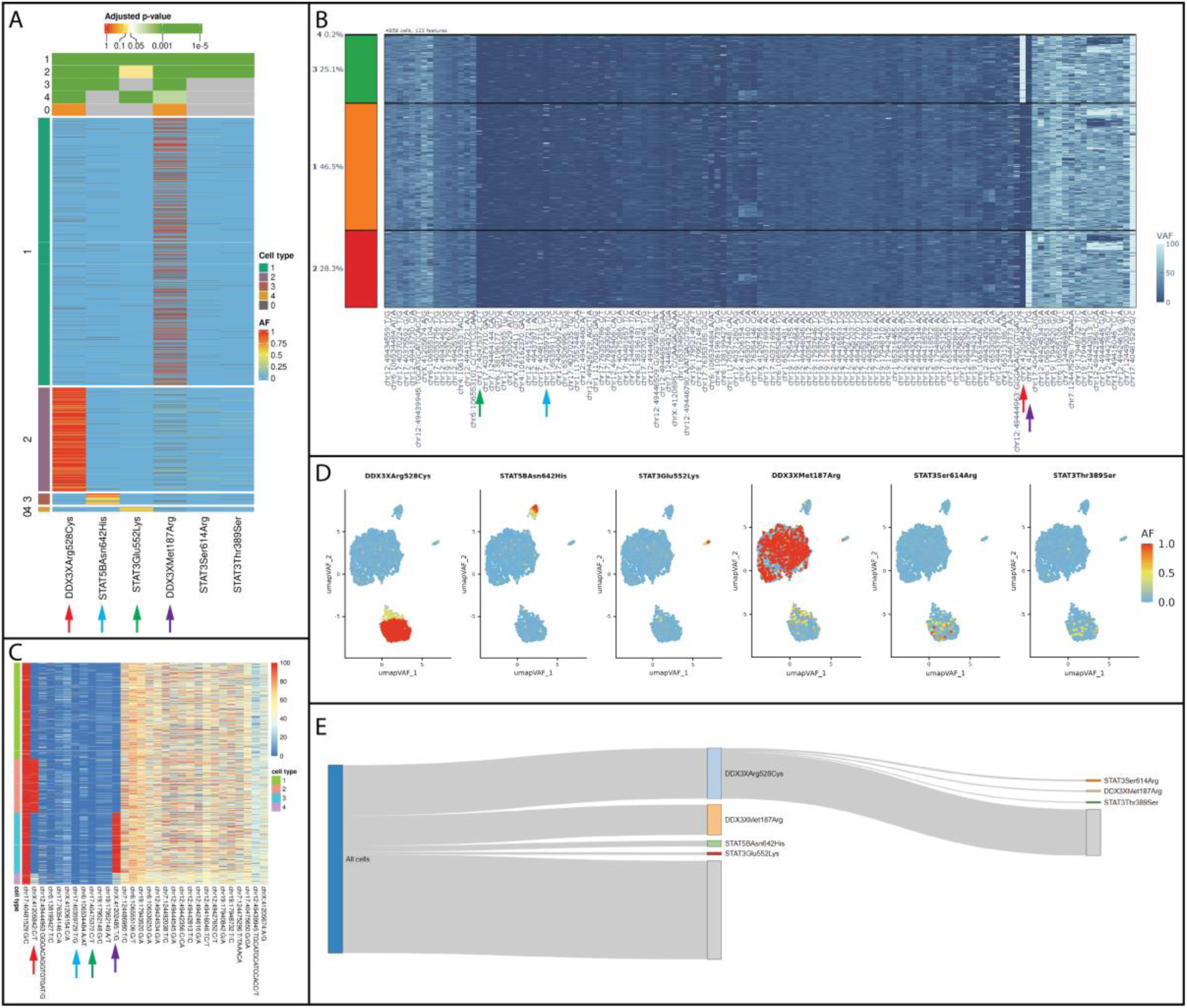
**Variant-based clustering and somatic cell type analysis in patient with refractory coeliac disease (RCD) using SCARCE, Optima and Mosaic**. A) Significant variants (p < 0.05, OR > 5) identified using SCARCE. Variant clustering and visualisation was also performed using Optima (B) and Mosaic (C). SCARCE visualisations further show localisation of allele frequency (AF) of enriched variants in cell sub-populations on UMAP (D) and limited co-occurrence (E).

A key feature of SCARCE is the ability to utilise ADT data from Tapestri sequencing to cluster and annotate cell populations, allowing users to identify somatic variant enrichment in a known cell type. SCARCE assists users in cell-type annotation by performing semi-automated annotation using the SCINA R package (Zhang et al. 2019) and producing plots which help the user decide on a final assignment for each cluster. Examples of these plots are shown in **Supplementary Figure 1**.

SCARCE is next run using ADT clustering for the same RCD patient analysed previously (**Figure 5**).

**Figure 5:**
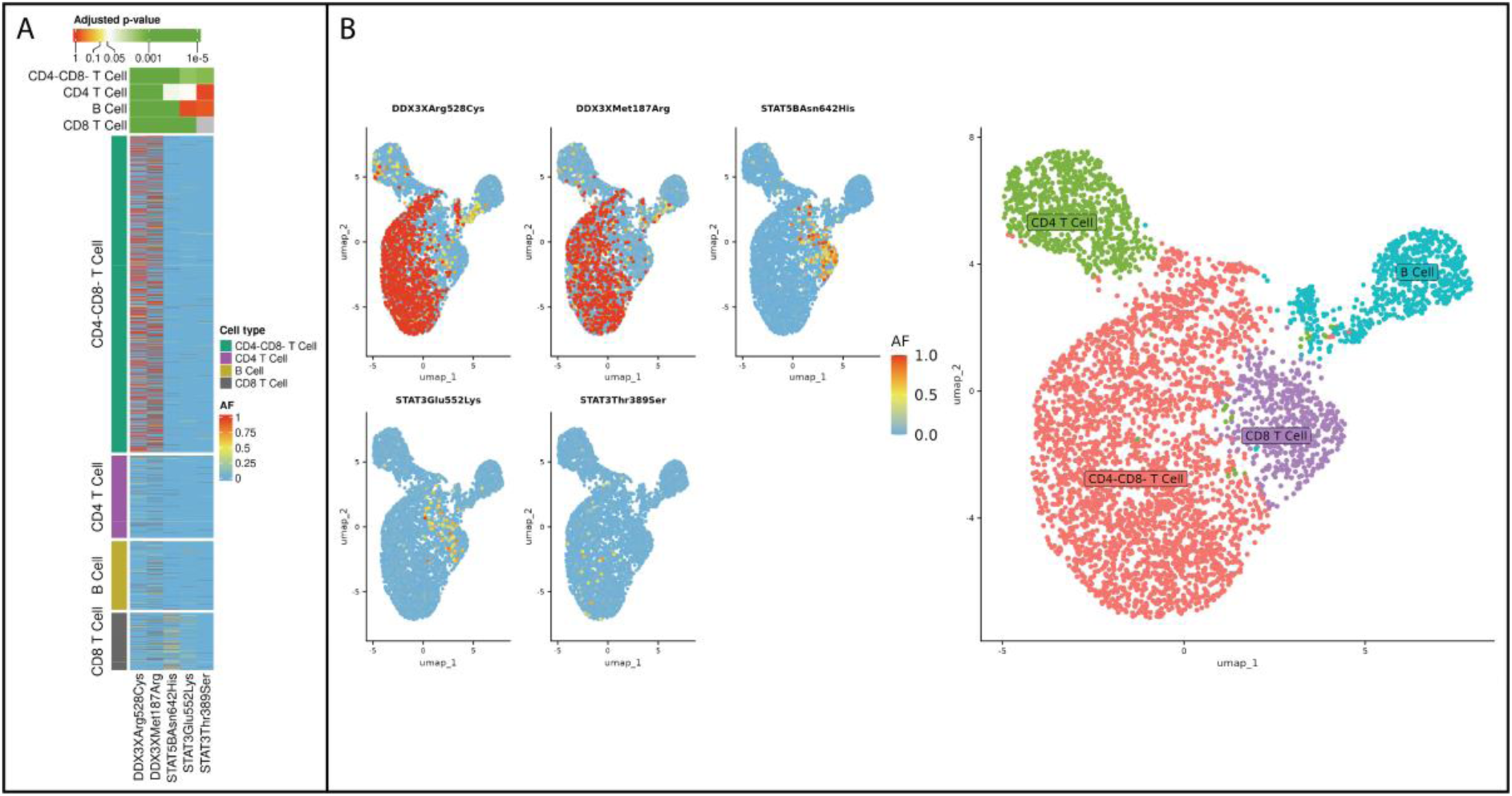
**SCARCE variant enrichment analysis using ADT surface marker-based clustering**. A) Heatmap of significant (p < 0.05, OR > 5) somatic variants identified by SCARCE. B) UMAP plot indicating allele frequency (AF) and localisation of significant variants.

Importantly, this workflow identifies the same known driver mutations as before, but reveals their cell-type enrichment. For example, both *DDX3X* driver mutations were found to be highly enriched in CD4-CD8-T cells, while *STAT5B^p.N642H^* and *STAT3^p.E552K^* were enriched in CD8+ T cells. These findings were, again, in agreement with the previous publication (Singh et al. 2025).

Finally, while SCARCE was designed to utilise the powerful multi-omic data capabilities offered by Tapestri or similar technologies, it can support any single-cell DNA sequencing technology that generates a multi-sample VCF where each ‘sample’ references a clone. To demonstrate this functionality, we used SCARCE to identify enriched variants in a patient with cryoglobulinaemic vasculitis who underwent clonally-amplified WGS performed on two aberrant cell populations (“blue branch” n=16, “orange branch” n=16, control n=2) (Singh et al. 2020b). Notably, these branches are known to differ by the presence of a *KLHL6^Leu90Pro^* variant, present only in the “blue branch” cells. Due to the vast amount of data available in WGS as opposed to the targeted sequencing performed by Tapestri, we limited this analysis to only chromosome 3 for this demonstration. **Figures 6A-B** show the results of this analysis. As expected, due to multiple hypothesis corrections across such a large number of variants and a small sample size in this study, no variants reached statistical significance. However, in such cases, users can instead focus on effect size. To do so, SCARCE offers the ability to filter results by Wald-Z score and OR instead, with the heatmap customisable to display these features. Doing so identifies *KLHL6^Leu90Pro^* among the top variants, along with several other previously unreported variants in this sample. Notably, the Sankey plot (**Figure 6C**) also highlights co-occurrence of many variants.

**Figure 6:**
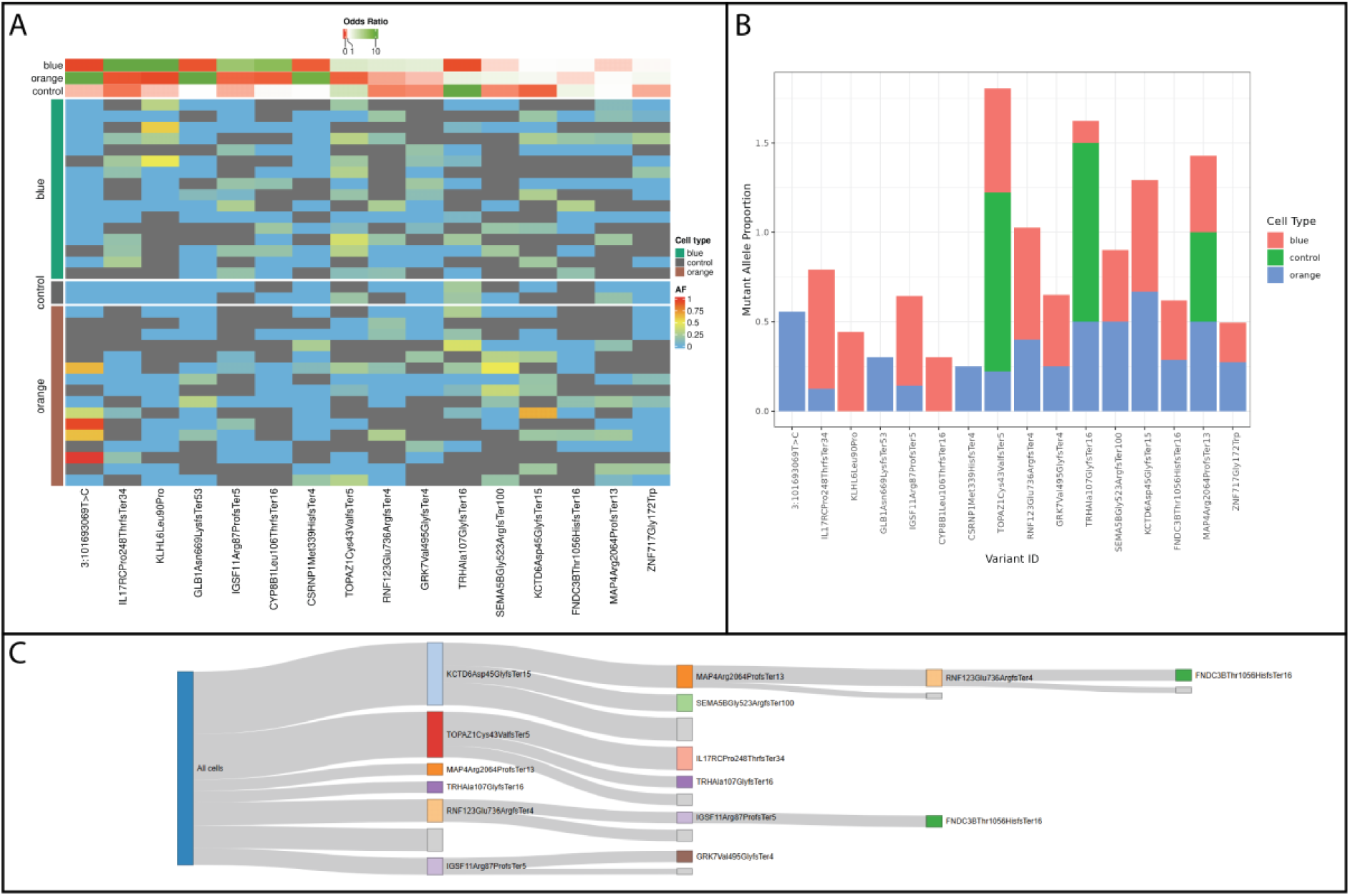
**SCARCE somatic variant prioritisation in clonally-amplified WGS sequencing**. Data consists of WGS performed on 34 cells from known clonal branches (blue n=16, orange n=16, control n=2). This analysis was restricted to chromosome 3 for demonstration purposes. SCARCE was used to generate a heatmap (A) of the top priority variants ordered by OR, as well as a bar plot showing mutant allele proportions per cell annotation (B) and a Sankey plot showing co-occurrence of variants (minimum set size =2) (C).

## Discussion

Here we present a simple and intuitive R package, SCARCE, to statistically prioritise and visualise somatic variants enriched in subpopulations detectable in scDNA-Seq data. Somatic mutations occurring in small clonally expanded cell populations are now understood to be key drivers in both cancer and autoimmune disease (Singh et al. 2020b; Singh et al. 2025), and can be extremely difficult to detect using traditional bioinformatics pipelines, reinforcing the need for new tools to be developed.

Recent years have seen rapid expansion of scDNA-seq library preparation technologies. For example, SCARCE works natively with HDF5 data from Tapestri, a technology which uses a microfluidic approach to capture single cells and perform targeted DNA sequencing of thousands of cells paired with ADT assays for surface marker expression (Pellegrino et al. 2018). However, SCARCE can be applied to any technology that produces multi-sample VCF-style files with one ‘sample’ column per cell. In this paper we additionally analysed clonally amplified data from isolated single cells using multiple displacement amplification (MDA), a precursor to NANOSeq, used by BioSkyb’s ResolveOME. ResolveOME employs whole-genome primary-template directed amplification (PTA) and sequencing with paired transcriptome sequencing of up to 384 isolated clones (Marks et al. 2023). This approach, therefore, sacrifices cell throughput but offers much greater sequencing capacity. Finally, we have tested on G&Tseq, an earlier technology which also offers paired genomic and transcriptomic sequencing of isolated single cells using MDA (Macaulay et al. 2015). Although these methods lack Tapestri’s ADT-based surface marker annotations, this can be recovered through flow cytometry-based isolation and annotation of individual cells prior to library preparation.

Despite the increasing number of single-cell DNA-based technologies, pipelines for somatic variant calling remain tailored to bulk sequencing approaches. Common tools like Mutect2 (McKenna et al. 2010), Strelka2 (Kim et al. 2018), and VarScan (Koboldt et al. 2012) can, in theory, be applied to single-cell data by considering each cell as a sample. However, these tools expect input of tumour and normal samples, while for single-cell data it is typically unclear which cells harbour pathogenic somatic mutations. SCARCE instead applies a 1 vs all approach to compare mutant allele counts between a specified cell population and all other cell types iteratively, statistically testing for enrichment in a cell population while prioritising deleterious variants based on Ensembl VEP annotations.

In this publication we compared SCARCE with MissionBio’s specific software Optima and Mosaic, which are intended for filtering and visualisation rather than statistical enrichment. Because they lack a prioritisation approach, they rely on stringent filtering to reduce, in this example, ∼100,000 variant calls down to ∼20-50 for clustering and visualisation purposes. SCARCE instead applies less-stringent filters to filter out only technical noise, followed by statistical enrichment to focus on probable somatic variants, an approach which reduces the risk of removing true positives. As demonstrated, SCARCE identifies cell-type enriched variants that are filtered out by other tools.

Another key feature of SCARCE is the ability to cluster cells using either ADT-or variant-based information. Comparing the two approaches on the same samples, we found that variant-based clustering is often better at isolating true rogue cell populations, as demonstrated by comparing **Figures 4C and 5C**. While variant-based clustering is also offered by Optima and Mosaic, our use of a greater number of prioritised variant features led to an improved ability to isolate populations, evident in **Figure 2**. On the other hand, ADT-based clustering allows SCARCE to accurately annotate cell types, giving users biological context for enriched somatic variants. For example, **Figure 5B** clearly and intuitively demonstrates biological cell-type specific enrichment of variants.

Single-cell sequencing data can quickly become extremely computationally expensive to analyse, depending on the number of cells analysed. We tracked the peak memory usage by each tool when analysing the two Tapestri datasets presented, and found that peak memory utilisation by SCARCE was far lower than Optima when analysing large datasets, thanks primarily to our use of chunked loading, though memory use was still much greater than Mosaic (**Supplementary Figure 2**). This means that moderately sized datasets (<5000 cells) can reasonably be run on a local machine without the need for high performance computing infrastructure.

Despite the advantages offered by SCARCE, we note several limitations which we aim to address in the future. For example, in its current implementation, SCARCE does not consider any form of structural variants, including copy number alterations, in its variant prioritisation. Additionally, completely automated cell-type annotation is not offered, requiring users to make informed annotation decisions based on the intuitive visualisations produced by the software. Finally, although SCARCE is designed to be memory efficient by implementing chunked-loading of variant data (which can be user-controlled), the software has been optimised for targeted sequencing panels rather than WGS. This means that analysing WGS data may incur higher than expected memory requirements and runtimes, an issue we plan to address in future releases.

Given the rapid expansion of scDNA-seq technologies, SCARCE’s development will continue in the future. In particular, there are several ways that cell clustering could be improved to better isolate cell populations of interest. For example, we have demonstrated that variant-based and ADT-based clustering both offer key advantages and have value independently for grouping cells. However, we expect that a method which combines both may be of significant value, potentially giving both better resolution for clustering and increased biological insight into enrichment results. Furthermore, while some technologies such as G&Tseq and ResolveOME provide paired genomic and transcriptomic data, SCARCE does not currently support cell clustering based on transcriptome data. Finally, in the specific context of autoimmune disease, where immunological clones can be typed based on repertoire sequencing of T-cell and B-cell receptors, we suspect that the ability to annotate clusters based on repertoire sequencing could offer improved resolution for identifying somatic variants in aberrant, self-reactive clones.

In summary, SCARCE offers users a powerful but simple way to prioritise somatic variants in heterogeneous scDNA-seq data. As scDNA-seq technologies advance, we aim to continue development of SCARCE to assist in the development of diagnostics and treatment options for patients with somatic mutation-driven diseases, including cancer and autoimmune disease.

## Methods

### Workflow overview

SCARCE allows two primary workflows and several optional steps dependent on data availability and user-preference (**Figure 1**). The ADT-clustering workflow is designed for experiments that include single-cell variant calls as well as surface marker expression from ADT assays. This workflow begins with semi-automated cell type annotation, which users can skip if utilising their own cell annotations, followed by enrichment of cell type-specific mutations and visualisation. In the variant-only workflow, SCARCE performs cell clustering based on present variants to identify rogue subpopulations of cells. These subpopulations are then annotated and investigated for variant enrichment as per the standard workflow.

### Input requirements

For the ADT-clustering workflow, SCARCE expects surface marker expression in the HDF5 file format, as supplied by Tapestri sequencing, or this can be skipped by providing user-defined cell annotations. Variant data can be provided in the same HDF5 object, or as a multi-sample VCF file with one column per cell. For the variant-only workflow, only the variant information is required, supplied as either an HDF5 object or multi-sample VCF.

### Data Preparation

Surface marker expression data are prepared using the *SCARCE::prepare_annotation_data()* function. ADT data is extracted from the HDF5 object and loaded into a Seurat (v5) object using *Seurat::CreateSeuratObject()*. The ADT matrix is normalised with the *Seurat::NormalizeData(normalization.method=”CLR”)* function and scaled using *Seurat::ScaleData().* Surface markers are filtered to a top user-defined percentile (default = 75%). Finally, PCA and UMAP dimensionality reduction are performed, with user-defined distance metrics (default = “euclidean”), and cell clusters are identified using *Seurat::FindClusters()* at user-defined resolutions (default = 0.2, 0.35, 0.5, 0.8, 1, 1.2).

### Custom Cell Annotation

SCARCE performs semi-automated cell annotation with *SCARCE::annotate_cells()* by first performing an automated per-cell annotation using SCINA (Zhang et al. 2019) and providing a series of plots to help the user ultimately decide on a manual per-cluster annotation (**Supplementary Figure 1**). By default, the CellMarker2.0 database provides a list of cell-surface markers associated with each cell-type, but these may be overwritten with user-defined markers. SCINA is run for a maximum of 100 iterations and assigns a single cell type and probability score to each cell. Cells with an assignment probability below a user-defined threshold (default 0.5) are annotated as “unknown”. Plots are produced informing the user of the surface marker expression and the SCINA cell type assignment proportions for each cluster. Finally, the user annotates cell clusters using *SCARCE::annotate_clusters*().

### Variant Processing and Filtering

Variant data are processed and filtered using *SCARCE::find_somatic_variants()*. Variant formatting, regardless of HDF5 or VCF input, is first normalised to a format that supports Ensembl Variant-Effect Predictor (VEP) input, and multi-allelic sites are expanded to become biallelic. Incorrectly reported variants that cannot be normalised are removed.

Variants are further filtered in three stages:

1. Call-level filters are used to filter variant calls for individual cells. In these cases, the numeric genotype (NGT) call is changed to 3.

- Depth < min_DP_per_call (default = 10)
- Genotype Quality < min_GQ_per_call (default = 30)
2. Variant-level filters are used to filter variant calls across all cells

- Alternate allele count < min_varcount_total (default = 30)
- Alternate allele proportion < min_varportion_total (default = 0)
- Alternate allele proportion > max_varportion_total (default = 0.7)
- Total number of cells with non-missing data < min_datacount_total (default = 100)
- Total proportion of cells with non-missing data < min_dataportion_total (default = 0.25)
- Minimum mean genotype quality < min_quality_mean_total (default = 0.3)
- Minimum mean allele frequency < min_allelefreq_mean_total (default = 0)
- Minimum mean allele frequency in cells classified as mutant < min_allelefreq_mean_mutants (default = 0.1)
3. Cell type-specific filters are used to filter variants in the cell type-specific output files

- Alternate allele count in cell type < min_varcount_celltype (default = 1)
- Alternate allele proportion in cell type < min_varportion_celltype (default = 0)
- Cell type cells with non-missing data < min_datacount_celltype (default = 1)
- Proportion of total mutated cells belonging to cell type < min_varportion_in_celltype (default = 0)

To help identify candidate driver mutations from the large number of spurious variant calls, variant annotations are generated using VEP (McLaren et al. 2016) including variant types, gene symbols, protein alterations, population allele frequency, and pathogenicity predictions using SIFT (Sim et al. 2012) and PolyPhen (Adzhubei et al. 2010). The –flag_pick_allele flag is used to ensure that a single annotation is returned per allele.

Based on VEP-annotations, a PRIORITY flag is assigned to variants with a maximum population allele frequency (--max_af) less than a threshold (default = 0.02) and their VEP consequence assigned as: missense_variant, stop_gained, stop_lost, start_lost, frameshift_variant, inframe_deletion/insertion or protein_altering_variant.

### Variant-Based Cell Clustering

Cells may be clustered and a Seurat object generated using the *SCARCE::cluster_variants()* function, similar to the method employed by Optima (Pei et al. 2023) and Mosaic (Zhang et al. 2023). This function first searches for the top n_features (default = 100) to use for clustering. Variants are first filtered to those with an assigned genotype in > 80% of cells. The Bernoulli variance (var_ngt) of the population alternate allele frequency (𝑝_alt_) is calculated as follows:

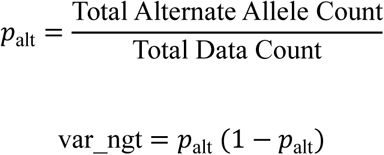

This down-weights both very rare and very high frequency (likely germline) variants. To retain informative very-rare events, we additionally select a set of rare variants (<2% frequency) restricted to mean genotype quality > 30 and rank these by descending alternate allele count. The final selection of n_features receives 90% from the top Bernoulli variance rankings and 10% from the rare variant rankings.

Using an allele frequency matrix of the selected variant features (𝑣) for all cells, we generate a cosine similarity matrix for all cells. For two cells, 𝑖 and 𝑗, the cosine similarity (𝑐𝑜𝑠) and distance (𝐷) are calculated using their allele frequency vectors across the selected variants (𝑥_𝑣_) only for the variants in which an allele frequency is available in both cells (𝑣 ∈ Ω_𝑖𝑗_):

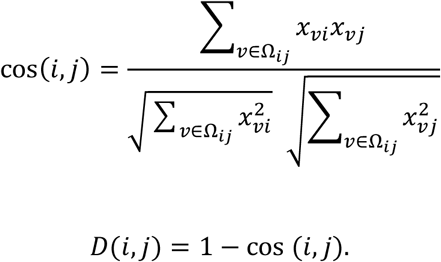

In cases where there is too small an overlap of data availability between cells (Ω_𝑖𝑗_ < 10), cosine distance is set to 1. This distance matrix is then embedded into 2D space using *uwot::umap*(n_neighbors = 30, min_dist = 0.3, metric = “euclidean”) and density-based clustering is performed using DBSCAN.

### Mutant Cell Annotation

*SCARCE::annotate_mutants()* is used to annotate mutant or diseased cell populations. This function can take a vector of variants of interest as input along with a Seurat object, and annotate cells carrying these variants as mutants, either taken as the intersection (those carrying all input variants) or union (those carrying any of the input variants) of cells. After doing so, *SCARCE::find_somatic_variants()* can be used to attempt to identify clonal variants present across this cell population.

### Cell Type Variant Enrichment

*SCARCE::find_somatic_variants(run_cell_type_enrichment=TRUE)* reports several statistics to help identify variants enriched in a cell type as specified by the provided cell type annotations. These annotations may refer to actual cell types as identified by surface protein expression using *SCARCE::annotate_clusters()*, variant clusters identified using *SCARCE::cluster_variants()*, or mutant cells identified using *SCARCE::annotate_mutants()*. Statistics are calculated by comparing each cell type of interest against all others in a 1 vs all approach, which is performed for all cell types (default) or only a user-defined selection.

For a given cell type, 𝐶, we collect the cell type-specific barcodes, 𝐵_𝐶_, and the barcodes for all other cell types 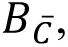 and compute:

- 𝑎: The number of cell type cells carrying a mutant allele (𝑁𝐺𝑇_𝐵𝐶_ ∈ {1,2})
- 𝑏: The number of cell type cells homozygous for the reference allele (𝑁𝐺𝑇_𝐵𝐶_ = 0)
- 𝑐: The number of “other” cells carrying a mutant allele 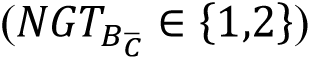
- 𝑑: The number of “other” cells homozygous for the reference allele 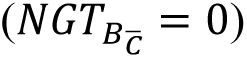

This yields a 2×2 confusion matrix per variant:

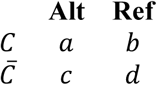

If any cell count is equal to 0, a Haldane-Anscombe correction is performed by adding 0.5 to each cell. The OR comparing 𝐶 and 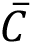 is:

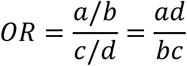

The standard error of the log odds-ratio is calculated as:

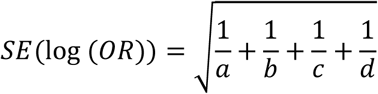

This is then used to calculate a Wald 𝑍-score for the log(OR):

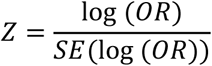

The Wald 𝑍-score is finally used to calculate a p-value on a standard normal distribution. For any case where min (𝑎, 𝑏, 𝑐, 𝑑) < 5, a two-sided Fisher test is used instead from *stats::fisher.test*. P-value adjustment is performed on p-values from all passing variants (FILTER=”.”) using the Benjamini Hochberg procedure.

### Reporting

In total, four key report files are generated by the *SCARCE::find_somatic_variants()* function and utilised by other downstream functions.

- Summary file (summary*.tsv): Contains one row per variant with VEP annotations, and variant-level summary information. Separate files are generated for all variants (summary.tsv), pass-only variants (summary_pass_only.tsv), and passing priority variants (summary_pass_only_priority).
- Cell type variant enrichment files (*cell_type*.tsv): Contains all the same information as the respective summary files with additional cell type-specific summary information and statistics for variant enrichment. Separate files are generated for all variants (summary.tsv), pass-only variants (summary_pass_only.tsv), and passing priority variants (summary_pass_only_priority).
- Genotype files: Contains all the same information as the summary.tsv file, with an additional column for each cell encoding per-cell allele frequency (AF), depth (DP), genotype quality (GQ), and numeric genotype (NGT) formatted as AF;DP;GQ;NGT.
- Variant counts per cell type: Contains all the same information as the summary.tsv file, with additional columns for each cell type containing the number of cells carrying a mutant allele and the total number of cells with data available.

### Visualisation

All plots are generated using the *SCARCE::plot_variants()* function. This function generates summary plots for a small number of variants of interest, which can be acquired by either retrieving the top N variants from the variant enrichment statistics, or by searching for a gene or specific variants of interest using the *search* and *search_col* arguments. The plots generated by this function include:

- Stacked bar plot: Plots the mutant allele proportions for each cell type
- Upset plot: Shows overlap and uniqueness of variants present between different cell types
- Allele frequency heatmap: A heatmap of the mutant allele frequency per cell clustered into their cell types
- UMAP plots: UMAP plots coloured by NGT and allele frequency
- Sankey plot: A Sankey plot showing the subclustering of variants in order from highest frequency to lowest

### Datasets

Tapestri sequencing data were retrieved from the Mission Bio data portal (portal.missionbio.com) for “patient 1” for AML single-cell measurable residual disease (scMRD) in HDF5 format.

The original data used in this publication, including the Tapestri data and the clonally-amplified WGS data were acquired as previously described (available under ENA: PRJEB31688 and NCBI PRJNA1088187) (Singh et al. 2020b; Singh et al. 2025).

### Software Availability

The SCARCE R package is available for download and installation from https://github.com/FieldLabFNQOmics/SCARCE

### Author contributions

SD, MS, CG and MAF conceived the project. SD, MAF, RL and FL performed preliminary analysis. RG developed the SCARCE package. All authors contributed to the manuscript, with most writing by RG, SD and MAF.

## ACKNOWLEDGMENTS

We thank the National Computational Infrastructure (NCI) for their computational support. The work was supported by National Health and Medical Research Council Fellowship APP5121190.

## SUPPLEMENTAL DATA

**Supplementary Figure S1:**
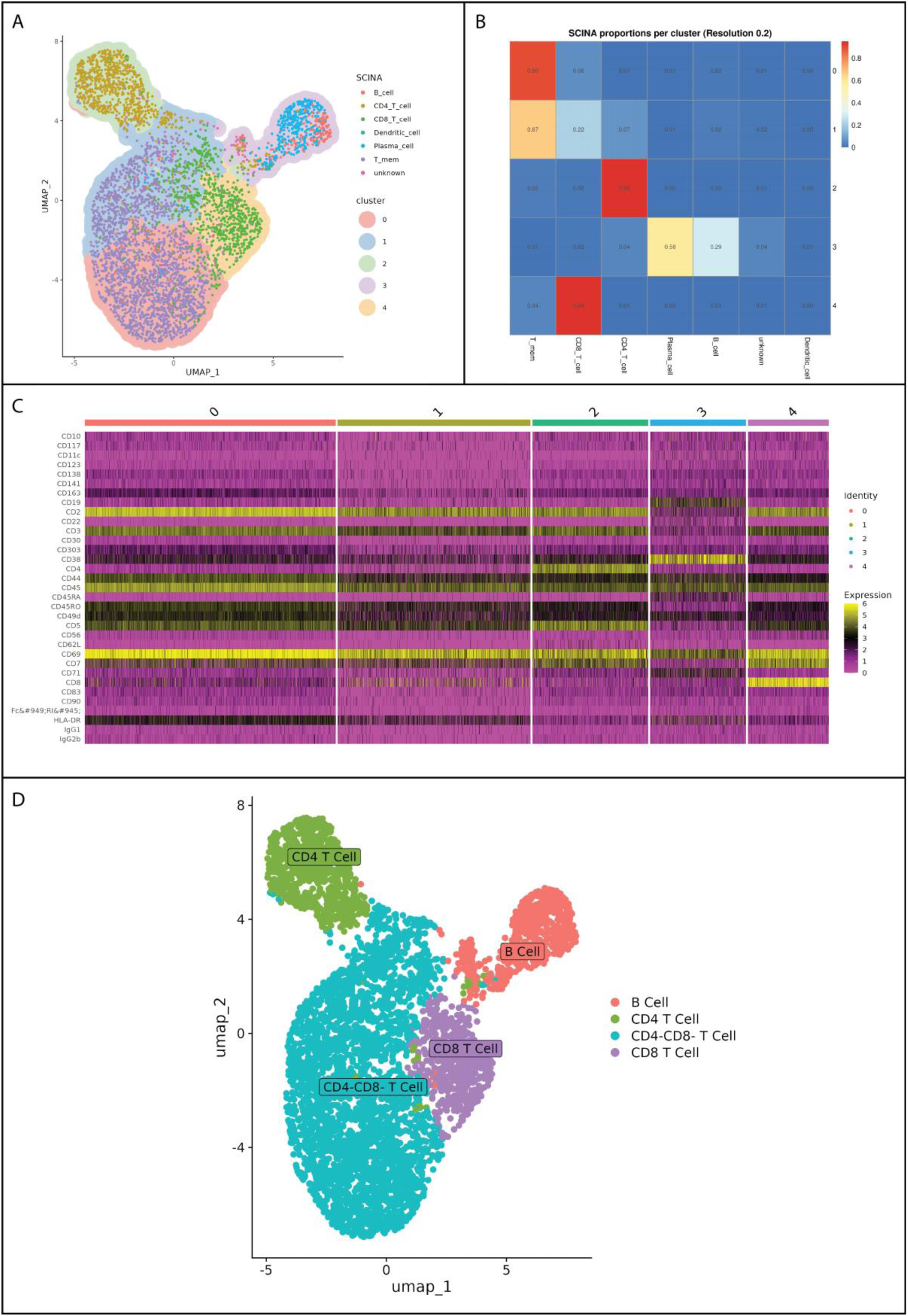
Semi-Automated Cell Type Annotation by SCARCE. The SCINA R package is used to automatically annotate a cell type to each individual cell based on known markers in the CellMarker2.0 database, or alternatively based on user-specified cell markers. A) A UMAP plot is produced which overlays the identified cell clusters with the SCINA-assigned cell annotations. B) A heatmap is produced to show the proportion of SCINA cell type assignments per cell cluster. C) A heatmap is produced which shows the expression of each cell surface marker grouped by cell cluster. Based on plots A-C, the user assigns a definitive cell type to each cluster, and a final UMAP plot is produced (D).

**Supplementary Figure S2:**
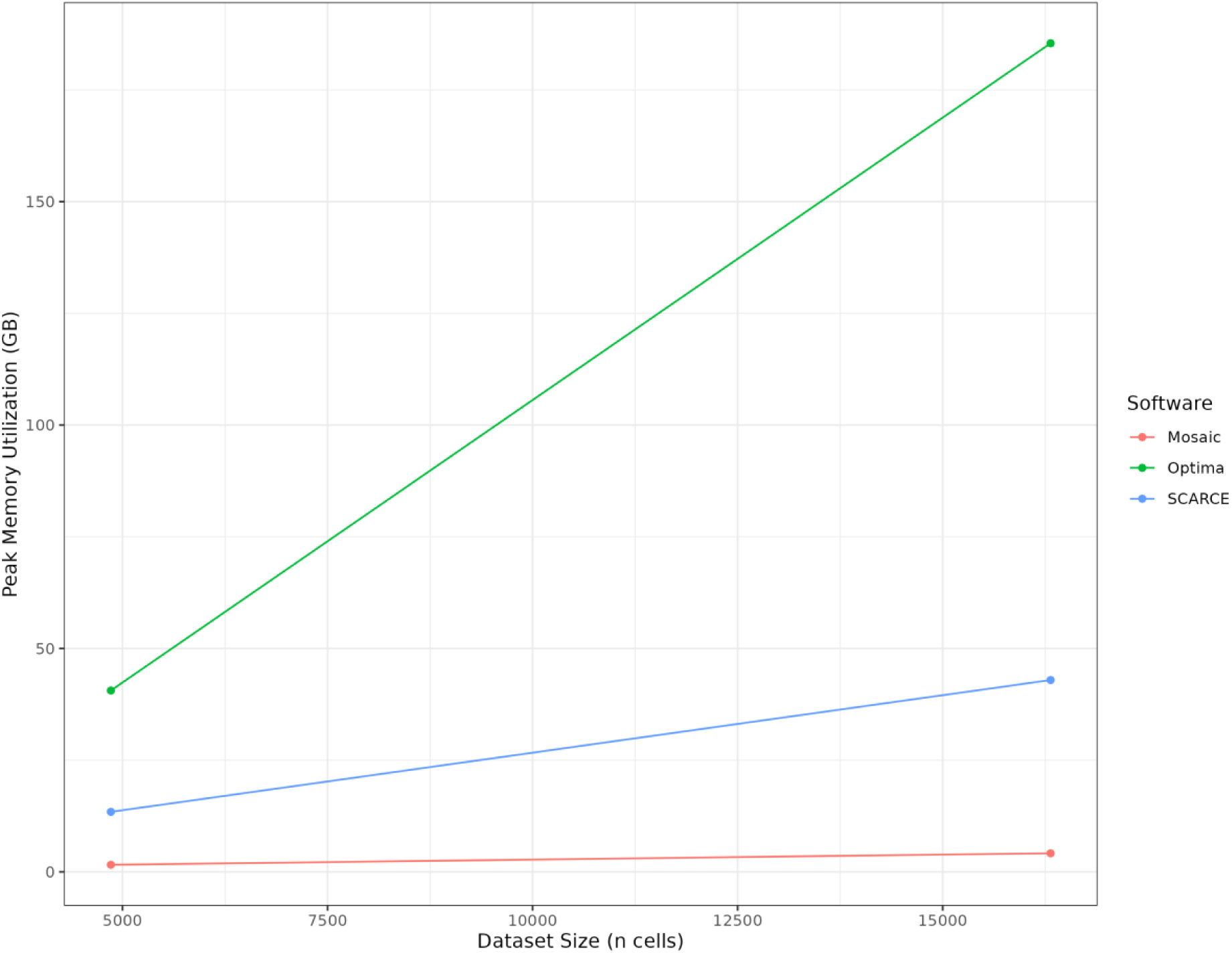
Peak Memory Usage Relative to Cell Number. Memory usage was tracked across for Mosaic, Optima and SCARCE across two MissionBio Tapestri data sets.

